# Hybrid approach for parameter estimation in agent-based models

**DOI:** 10.1101/175661

**Authors:** Jill A. Gallaher, Andrea Hawkins-Daarud, Susan C. Massey, Kristin R. Swanson, Alexander R.A. Anderson

## Abstract

Agent-based models are valuable in cancer research to show how different behaviors emerge from individual interactions between cells and their environment. However, calibrating such models can be difficult, especially if the parameters that govern the underlying interactions are hard to measure experimentally. Herein, we detail a new method to converge on parameter sets that fit an agent-based model to multiscale data using a model of glioblastoma as an example.

## I. Agent Based Models and Glioblastoma

Agent-based models (ABMs) are useful in cancer research to study evolutionary dynamics of cell populations and emergent behavior from cell-cell and cell-environment interactions [1]-[4]. As we gain a deeper understanding of the spatial and temporal heterogeneity in tumors [5], [6] and develop better techniques for measuring single cell dynamics [7], an ABM can be an invaluable tool to integrate these data and to understand the underlying mechanisms of tumor progression. However, ABMs often have many parameters, which may involve unmeasured rates and interactions. Here we describe a method to estimate a large set of model parameters, used to fit glioblastoma multiforme (GBM) model, to multiscale experimental data.

GBM is a particularly heterogeneous and malignant brain cancer, characterized by infiltration of individual cells deep into the brain tissue. The abundant invasion through essential brain anatomy and heterogeneity makes curative intervention difficult, if not impossible. To investigate the role of environmental factors involved, an experimental rat model was created using platelet-derived growth factor (PDGF) as a driver of tumor initiation and progression [8]-[10]. The experiment yielded multiscale data: bulk tumor size measurements (from imaging), cell population sizes (using fluorescence microscopy), and single cell data (by tracking individual cells *ex vivo*). Of particular interest, the single cell data revealed marked heterogeneity amongst cells, which we sought to capture using a hybrid ABM.

## II. GUIDE TO THE METHOD

While there are many parameter-fitting techniques that can be used for continuous models, issues arise with ABMs due to their stochastic nature, which potentially corresponds to large fluctuations in behavior from small parameter changes and even large variability in output from a single set of parameters [11]. Additionally, ABMs may have a large number of unknown parameters and simulation times may be very long. Simple methods like random sampling and parameter sweeping can be used to find a loose fit and get a sense of the overall system behavior when there are few unknown parameters. However, with more complex systems that have many unknown parameters, schemes such as evolutionary algorithms, Bayesian approximations, swarming methods, and simulated annealing should be used [11]-[14].

Our hybrid ABM has 16 unknown input parameters and 16 experimentally measured output metrics. After encountering poor convergence using random sampling, a genetic algorithm, and simulated annealing, we developed a hybrid approach that achieved a good fit in less time. Our method uses a high-performance computing cluster (HPCC) to iteratively narrow a large parameter space to obtain a set of solutions within an allowed error. Figure 1 shows an overview of the procedure. For each iteration, the parameter space is refined using both a genetic algorithm approach combined with random weighted sampling. With the genetic algorithm we use selection to directly inherit the best fits in the set and mutation to allow divergence, so that improvements are possible while also preserving the correlation between parameters. Random sampling from distributions generated from the winning parameters was used to search more globally and avoid local minima.

**Figure 1.**
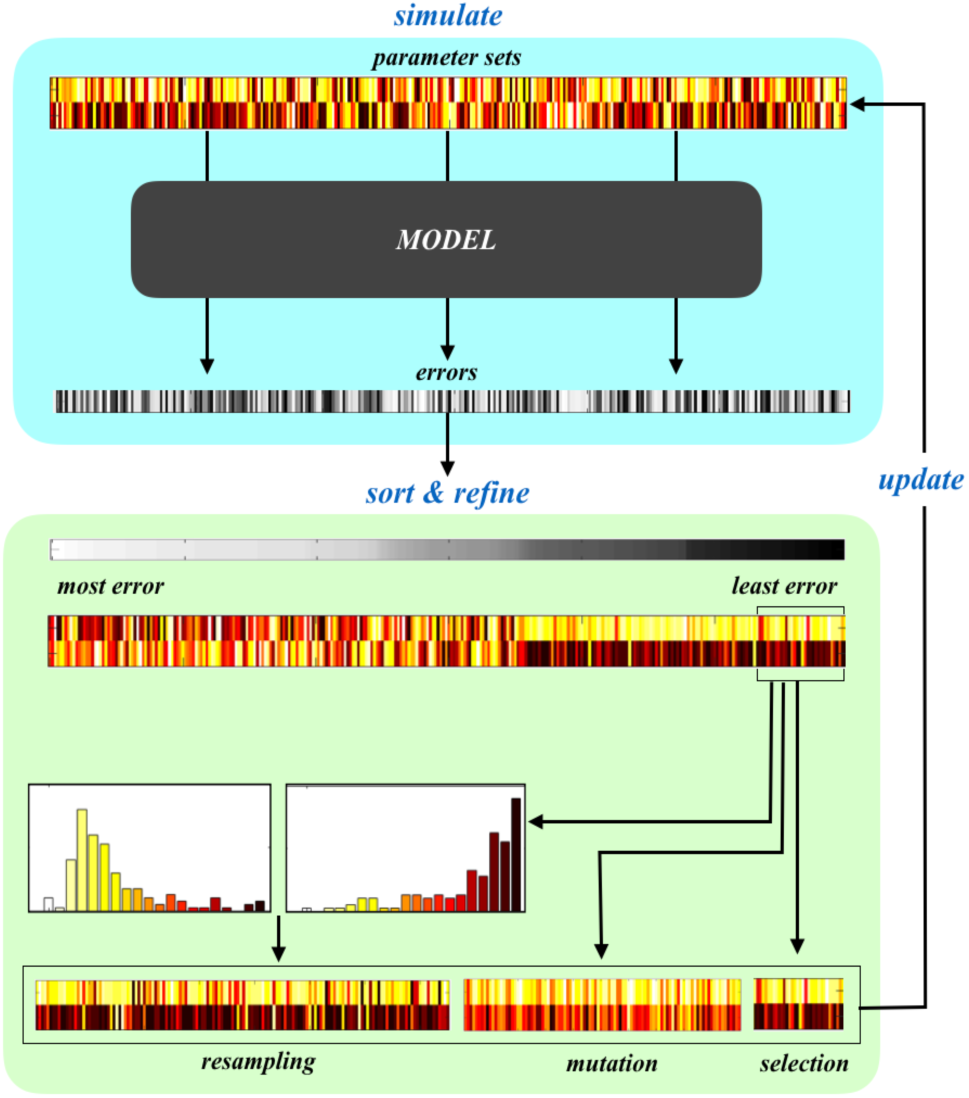
Illustration of method for a simple case of fitting two parameters to data. The parameter estimation procedure iteratively refines a large number of parameter vectors. At each generation, simulations, initialized with each parameter vector, are run in parallel, and the outputs are compared with the data to get a total error value. The errors are sorted to find the best fits, which are used to determine the next generation of parameter vectors. The selection method combines direct inheritance of the best fits (selection), random adjustment of the best fits (mutation), and random sampling from each parameter distribution generated from the best fits (resampling).

### A. Defining inputs and outputs

#### Parameter range selection

Each parameter range needs to be bounded within reasonable biological values. Given no prior knowledge on reasonable distributions, we uniformly sample the space to populate a large number of parameter vectors. In this example, we take 5000 samples, due to the size of the parameter space and the available computational resources[11], [13].

#### Mapping model outputs to experimental data

The ABM needs to be able to match its outputs to measures analogous to the experimental data. While this data set includes tumor sizes, ratios of different cell types, and single cell proliferation events and velocity distributions at various time points, we will only elaborate on how we transformed the single cell distributions into a measure of tumor size.

Extracting a single metric for tumor size using an ABM is not straightforward since the model tumor is a very diffuse and oddly shaped set of single cells (Figure 2). To calculate a radius, we first created a radial mesh from the center of the tumor mass and recorded the number of cells at each location. To find the total density at each location, we divided the cell number, *N*_*θ*_, by the total area:

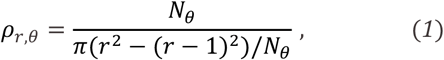

where *r* is the radial distance from the center point at r=0 and *N*_*θ*_ is the number of angular divisions defining the radial grid. In the experiment, the average diameter was found from measuring volumes of abnormal hyperintensity on magnetic resonance (MR) images. While there is no accepted mapping of tumor cell density to MR intensity, we used a cell density threshold of 50% for the model. The diameter was then found by averaging over all angles the maximum distance from the center of the tumor that 50% cell density was observed (Figure 2).

**Figure 2.**
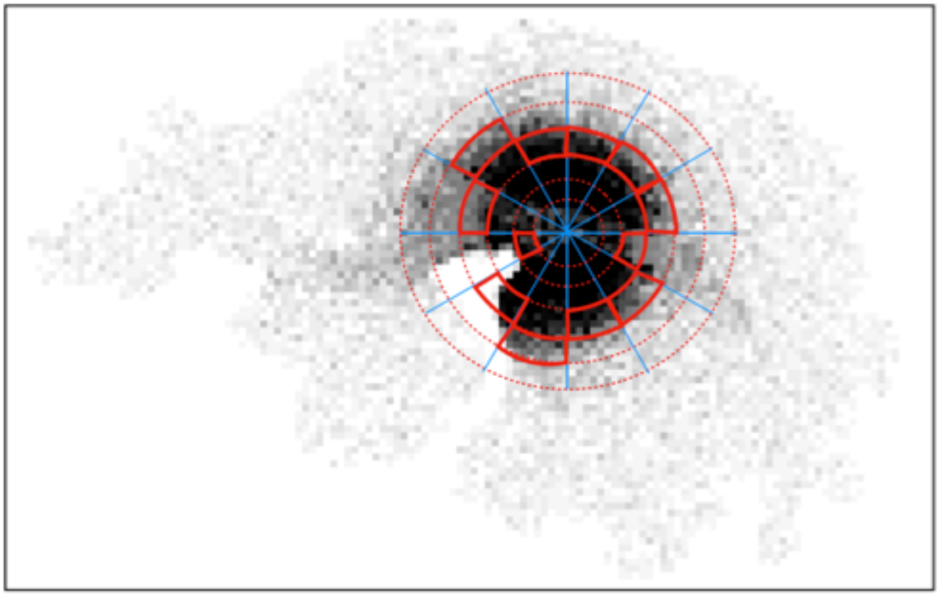
Mapping single cell densities to estimate average tumor diameter. The background shows cell density distributions on a regular mesh. To calculate the average tumor radius, single cells are mapped to a radial mesh where the tumor edge is defined as the largest radial distance for each angle with a density of ≥ 50% of the carrying capacity. The radius is found by averaging this distance over all angles.

### B. Parallel computation and error calculation

#### Running the model

The process of running a simulation for each parameter vector is perfectly parallel for each generation, meaning each simulation is independent of the others thus allowing them to run simultaneously. The run time per simulation was between 13 and 30 minutes. However, the scheduler on our HPCC limits users to 1000 simultaneous jobs, therefore a typical generation composed of 5000 runs took between 65 and 150 minutes.

#### Error Calculation

The model outputs that corresponded to data measurements were recorded during each simulation run. To get a total error from these outputs, the individual errors were weighted. This accounts for the allowed variance of each metric individually, in order to compare values of different metrics on different scales. For each output value *m*_*i*_, we calculate the error *E*_*i*_ from the data value *m*_data_, using a normalized weighted least squares formula:

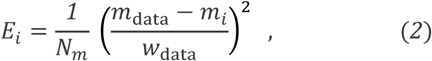

where *N*_*m*_ is the number of metrics, ensuring the sum of all errors 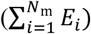 in a single run will be equal to or less than 1 if the values *E*_*i*_ on average are within the allotted weighted margin, *w*_data_, of each output.

### C. Sort errors to get best parameter combinations

After running the simulations on the HPCC, the selection of the next parameter set is done in MATLAB. From the output vectors, we sum the error vector for each run. We found that occasionally error would build up in a single metric, so with the constraint that a single error cannot exceed a threshold value, we ranked the total errors in ascending order with their corresponding parameter vectors. From the ranked set, we take the top 10% to create the next generation of parameter vectors.

### D. Refine the parameter sets

The winning parameter vectors are directly inherited and continue to the next generation, while the remaining 90% are found by mutation (40%) and resampling (50%). For mutation, each parameter was allowed to vary within a normal distribution centered at the original value with a standard deviation of 5% of the range. For resampling, a distribution was created for each parameter by binning the range into 20 equal intervals and recording the counts. Each parameter was sampled independently. We considered the parameter estimation procedure converged when the parameters of the current generation did not substantially change mean (by 5% of the range) or standard deviation (by 0.5% of the range) from the previous 2 generations. This system generally took around 10-12 generations to fully converge.

## III. DISCUSSION

Parameter fitting, model analysis, and data integration for ABMs is extremely difficult due to their stochastic nature. Randomness and heterogeneity amongst the many individual agents makes these models subject to noisy output, and the bottom-up approach of ABMs means that many underlying, unmeasured parameters need to be estimated. Random sampling and parameter sweeping are useful methods for investigating the global effect of parameter changes in an ABM. However, these methods are inappropriate when the number of unknown parameters is large. Finding optimal fits to data requires more complex, automated algorithms.

We are ultimately interested in characterizing heterogeneity in GBM on the single cell level to understand individual cell dynamics in disease progression and response to therapy. The noise generated from run to run due to the heterogeneity in single cell behavior, even with the same parameters could often be very large. We alleviated this concern by running a large number of parameter vectors in parallel over several generations. Some methods automate the entire process by using feedback during the simulation process [12], [15], which might improve parameter selection dynamically to decrease convergence time. The method presented here runs the simulations and refines the parameter array in separate steps, but also avoids complex cluster scheduling procedures and allows the user to observe the results at each step and make any necessary adjustments.

Using this approach we were able to converge on a set of parameters that fit bulk data (e.g. tumor size) and individual data (e.g. distributions of migration rates) to a reasonable degree. Beyond fitting the model to data, *it helped us better understand which parameters were critical and which ones had little influence on the measured output*. This method is a hybrid approach; so further refinements of each component are possible. However, this practical and easy to implement technique balances parameter correlation preservation with a more global search strategy to aid in systematically and efficiently narrowing a large parameter space. Whilst we used a GMB model here, our approach is equally applicable to any complex ABM coupled with appropriate multiscale data.

## ACKNOWLEDGMENT

We thank the organizers of the Maths of the CSBC and PSON meeting that inspired the creation of this handbook. This material is based upon work supported by the James S. McDonnell Foundation Collaborative Activity Award #220020264.

